# Fibroblast fusion to the muscle fiber regulates myotendinous junction formation

**DOI:** 10.1101/2020.07.20.213199

**Authors:** Wesal Yaseen-Badarneh, Ortal Kraft-Sheleg, Shelly Zaffryar-Eilot, Shay Melamed, Chengyi Sun, Douglas P. Millay, Peleg Hasson

## Abstract

Vertebrate muscles and tendons are derived from distinct embryonic origins yet they must interact in order to facilitate muscle contraction and body movements. How robust muscle tendon junctions (MTJs) form to be able to withstand contraction forces is still not understood. Using techniques at a single cell resolution we reexamined the classical view of distinct identities for the tissues composing the musculoskeletal system. We identified fibroblasts that have switched on a myogenic program and demonstrate these dual identity cells fuse into the developing muscle fibers along the MTJs facilitating the introduction of fibroblast-specific transcripts into the elongating myofibers. We suggest this mechanism resulting in a hybrid muscle fiber, primarily along the fiber tips, enables a smooth transition from muscle fiber characteristics towards tendon features essential for forming robust MTJs. We propose that dual characteristics of junctional cells could be a common mechanism for generating stable interactions between tissues throughout the musculoskeletal system.

---

Transplantation experiments in avian embryos as well as genetic investigations in mice performed over the last few decades identified the lineages that make up the vertebrate limb musculoskeletal system. These have demonstrated that while myogenic precursor cells are somite-derived, the connective tissues, tendons and bones are derived from the lateral plate mesoderm (LPM) ^1–4^. However, novel single cell resolution techniques allow us to revisit these early observations and focus on specific regions such as the sites of interaction between the distinct tissues to resolve their cellular contributions. One such focal point is the myotendinous junctions (MTJs).

Although being critical for muscle functioning in transmitting the force generated by the muscle to the tendons and skeletal elements, our understanding of the mechanisms that underlie MTJ development and maintenance are still relatively unclear. Importantly, while myofiber tips along the MTJs serve as the sites of interaction with the tendons, during embryonic and neonatal development when myofibers elongate via myoblasts’ fusion, the majority of fusion events also take place at these regions^5–9^. Accordingly multiple signaling pathways such as BMP and FGF signaling are tightly coordinated along the fiber tips ^10,11^ although their exact contribution to fusion, myofiber elongation and/or MTJ formation and maintenance is still unclear.

To examine the mechanisms underlying MTJ formation we have carried out a single cell transcriptome (scRNAseq) analysis of the MTJ region. This analysis revealed the presence of a unique cluster of cells expressing both myogenic as well as fibroblastic characteristics. Interestingly, cells in this cluster also express multiple genes known to be associated with MTJs. Surprisingly, this analysis further identified that the cells expressing the secreted extracellular matrix (ECM) modifying enzyme Lysyl oxidase-Like 3 (LoxL3), an essential enzyme required for MTJ formation, that is secreted from myofiber tips ^12^ is expressed by fibroblasts and not by myogenic cells. Using fluorescent in situ hybridization (FISH) analyses and tracing of fibroblast nuclei in vitro and in vivo within the developing muscle we identify a novel mechanism by which LPM-derived fibroblasts transdifferentiate, switch on myogenic characteristics and fuse into the myofibers along the MTJs. We suggest recruitment of LPM-derived cells from muscle boundaries and their fusion into the myofibers is essential for normal MTJ development ensuring proper localization of proteins along these junctions.

## scRNAseq reveals cells with dual identities

To investigate the cellular contributions to the MTJs, we carried out a single cell transcriptome analysis of the muscle-tendon region at P0. At this stage, MTJs have already formed yet myoblast fusion at this region is high. Single cells (i.e. without contribution of syncytial myofibers) were used for the analysis. Transcriptome analysis identified 21 independent clusters (Fig. 1a). Clusters expressing the pan muscle connective tissue interstitial marker *PDGFRα* (marked as fibroblasts in the UMAP; Fig. 1a) and *Osr1* were identified. These included muscle connective tissue fibroblasts with high *Tcf4* (*Tcf7l2*) or fibro-adipogenic precursors (FAPs) expressing *ScaI* (*Ly6a*). These fibrogenic clusters expressed significantly distinct transcriptomes from those enriched for myogenic markers (consisting of myoblasts progenitors and *Myogenin* expressing myocytes; Supp. Fig. 1a). We could further identify tenocytes, endothelial and mural cells (Supp. Fig. 1b-d). A small number of cells co-expressing myogenic and fibrogenic markers were identified in multiple clusters. Notably, while generally the above clusters could be associated with a single identity, a small number of cells clustered together (n=38; dual identity cells) expressing both fibrogenic identity markers as *PDGFRα, Osr1, Col1a1 and Dcn* but also myogenic markers such as *Pax7*, *Myf5*, *MyoD1*, *M-Cadherin* (*Cdh15*) and others (Fig. 1b-e and Supp. Fig. 1a-g). Interestingly, cells in this cluster also express multiple MTJ and tenogenic markers such as *Ankrd1, Thbs4, Tn-C and Bgn* ^13–16^ (Fig. 1e).

**Figure 1.**
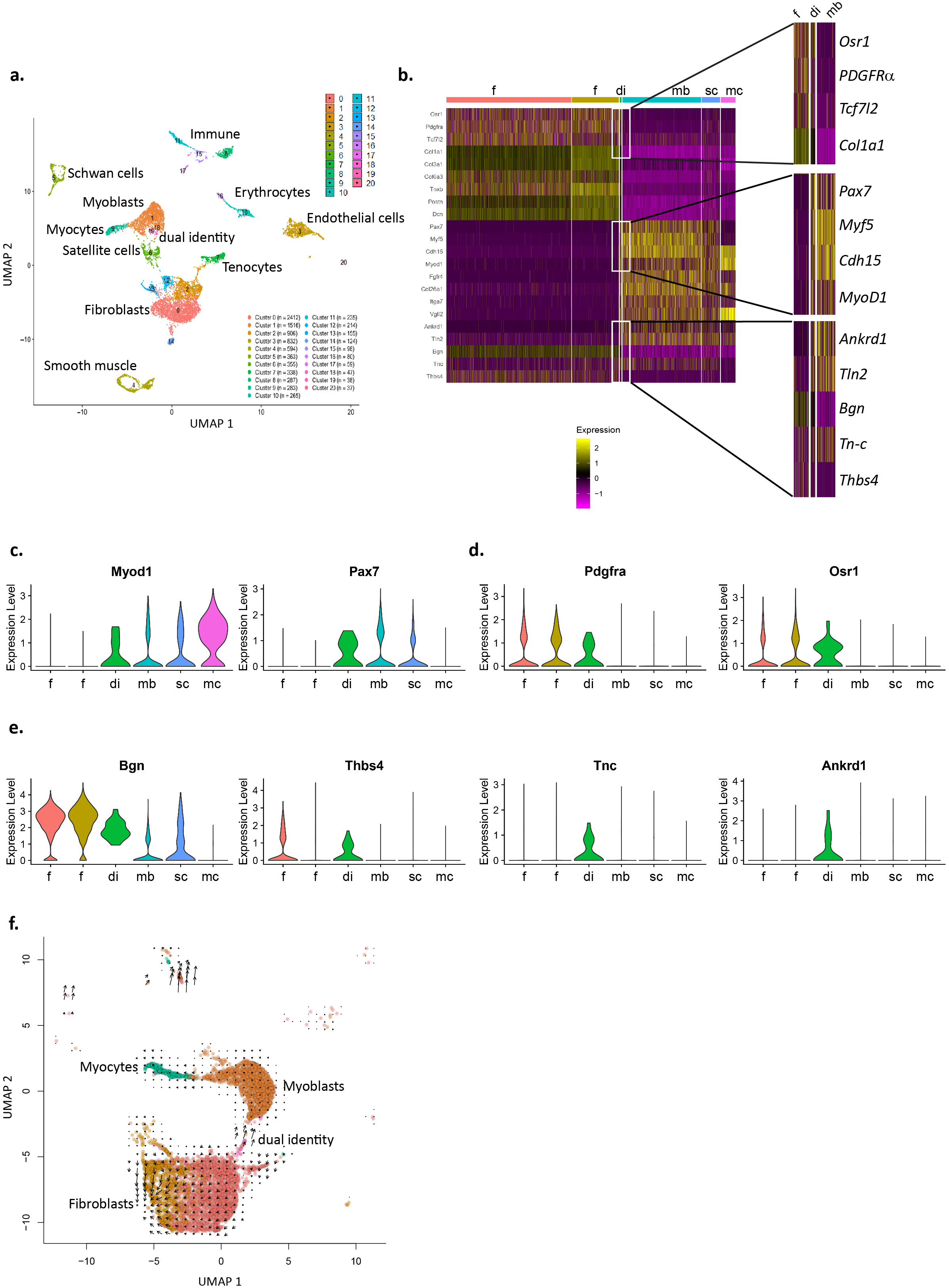
scRNA sequencing reveals cells with dual myogenic and interstitial transcriptional identities. UMAP depicting cell clusters (a) and violin plots of distinct genes associated with myogenesis (b) FAPs/fibroblasts (c) and tendon and MTJs (d). UMAP of RNA velocity analysis of myogenic (1,6,9), fibrogenic (0,2) and dual identity clusters (e). f - fibroblasts; mb - myoblasts; mc - myocytes; sc - satellite cells; di - dual identity.

To further characterize the dual identity cells, we used Ingenuity Pathway Analysis (IPA) software. When comparing transcriptomes of this cluster to that of the myogenic clusters using the upstream regulator analysis tool, we find TGFβ, previously associated with MTJ formation ^17^, being the most highly upregulated pathway in this cluster (Z score 5.528, p value 1*10^−37^). This analysis further suggests BMP signaling, known to be highly activated along myofiber tips ^11^, is activated in the dual identity cells (Z score 3.544, p value 5.7*10^−7^). TGFβ and BMP signaling were also suggested to be activated when comparing these unique progenitors to fibrogenic clusters although to a lower extent (Z score 1.69, p value 3*10^−10^; Z score 2.13, p value 2.95*10^−3^, respectively) altogether suggesting these two cascades are significantly activated in the cluster containing the dual identity cells.

The dual identity cells expressed genes associated with myogenic proliferation (e.g., *Pax7* and *Myf5*) and early differentiation (e.g., *MyoD1* but not *Myogenin*). Accordingly, IPA analysis suggests they are much more motile and proliferative. WEB-based Gene Set Analysis Toolkit (WebGestalt^18^) using the differentially expressed genes (DEGs) in this cluster versus those of the myogenic or fibrogenic clusters was further carried out. Changes in ECM composition of the dual identity clustered cells only vs. myogenic clusters were highlighted. These differences further demonstrate that cells in this unique cluster express high levels of ECM genes more similar to that observed in fibrogenic and tendon cells reinforcing the notion these cells form a transition between the myofiber and the tendon within the MTJ.

Remarkably, RNA velocity, an analysis which predicts future cell fate based on abundance of nascent (unspliced) and mature (spliced) mRNA forms in single cell data ^19^ suggests that the progenitors with a dual identity stem from fibrogenic clusters moving towards the myogenic identity (Fig. 1f and Supp. Fig. 1h). Hence raising the possibility that these progenitors transdifferentiate from fibroblasts into myogenic progenitors.

## *LoxL3* RNA is expressed by muscle interstitial cells

To begin to test the transdifferentiation idea, we probed a system where we previously demonstrated that the ECM modifying enzyme LoxL3 plays an essential role in MTJ development. In *LoxL3* mutant embryos, myofibers do not properly anchor along the MTJs ^12^ (Supp. Fig. 2a-c). The scRNAseq analysis revealed *LoxL3* transcripts are primarily expressed in fibrogenic clusters and not in the myogenic ones (Supp. Fig. 1i). Surprisingly, this expression pattern contrasts with the expression of LoxL3 protein which is expressed inside the myofiber tips (Supp. Fig. 2d-k) and being a secreted ECM-modifying enzyme is presumably secreted from their tips to modify the MTJ matrix promoting myofiber anchorage along the MTJ.

To verify the scRNAseq results that suggest *LoxL3* RNA is expressed primarily by fibroblasts we carried out FISH analysis on tissue sections. In accordance with the above RNA sequencing results and in contrast to LoxL3 protein expression, we find *LoxL3* RNA is mostly expressed adjacent to the muscle tips yet primarily outside of the myofibers (Fig. 2a,b).

**Figure 2.**
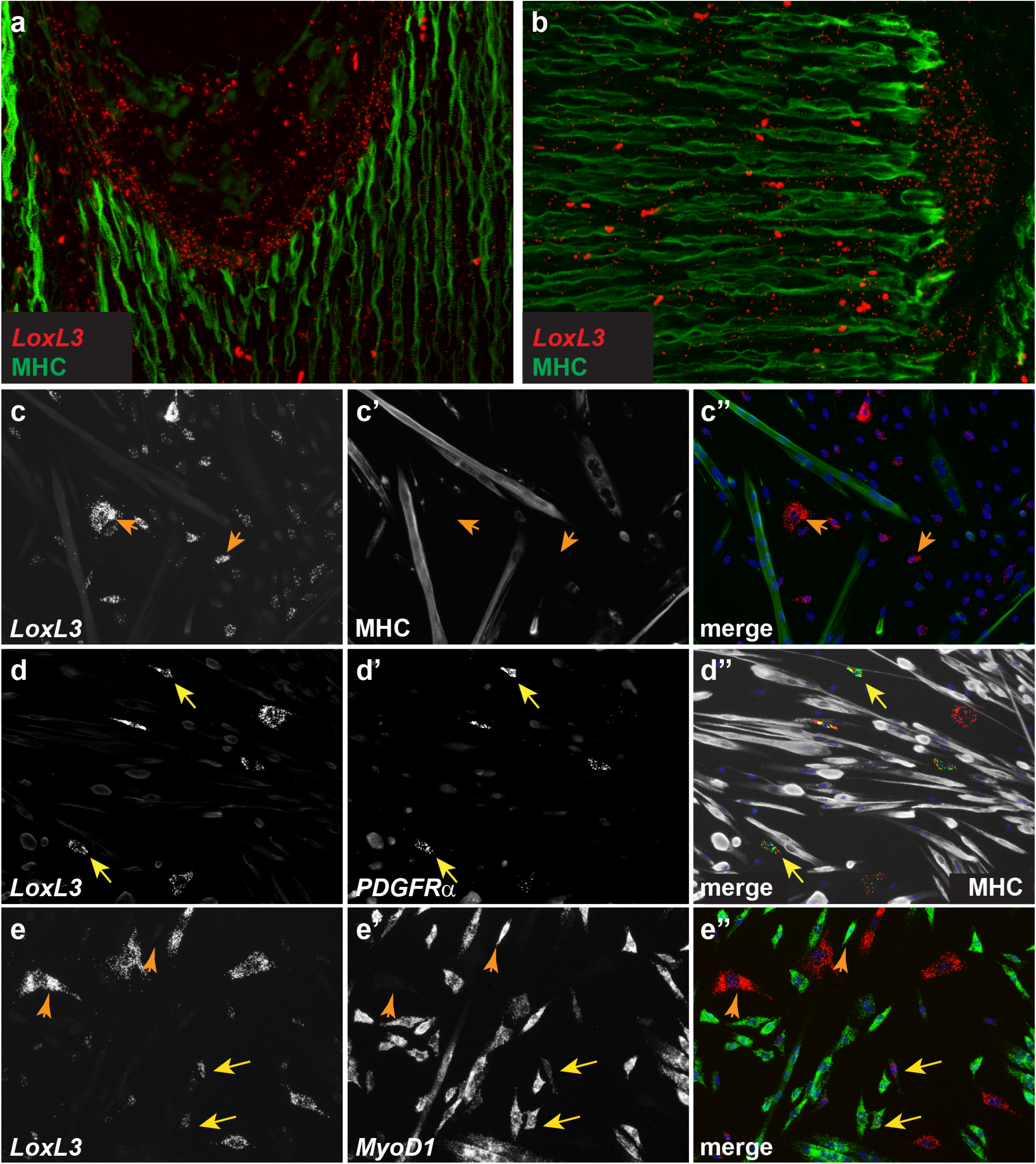
*LoxL3* RNA is expressed in muscle interstitial cells. FISH staining for LoxL3 RNA (red) and MHC immunostaining (green) at E15.5 demonstrates LoxL3 is expressed close to myofiber tips yet primarily outside of the myofibers (a,b). FISH for *LoxL3* (c) or also for *PDGFRα* (d) and anti-MHC staining demonstrates that *LoxL3* RNA is not expressed in myofibers but in *PDGFRα* expressing interstitial cells. FISH for *LoxL3* and *MyoD1* (e) reveals that most cells express either one of these markers (orange arrows), some express both (yellow arrows).

This unexpected observation of different protein and RNA expression lead us to further explore *LoxL3* RNA localization in a more controlled environment. We took advantage of our previous observations demonstrating that in in vitro differentiated myotubes, LoxL3 protein localization is maintained at the tips ^12^. We therefore monitored *LoxL3* RNA in primary cultures containing myoblasts, myotubes (marked by myosin heavy chain, MHC) and interstitial fibroblasts/FAPs (marked by *PDGFRα*, Supp. Fig. 2l) (Fig. 2c-e). In accordance with the in vivo results, no *LoxL3* RNA expression was observed in myotubes (Fig. 2c-c’’). In contrast, *LoxL3* mRNA was primarily observed in *PDGFRα* expressing fibroblasts/FAPs interstitial cells (Fig. 2d-d’’). Staining for *LoxL3* and *MyoD1* RNAs revealed that most cells expressed either marker - myogenic cells expressed *MyoD1* whereas fibroblasts/FAPs expressed *LoxL3* (Fig. 2e-e’’; orange arrowheads). Notably, ~33% of the *LoxL3* expressing cells also expressed *MyoD1* (Fig. 2e-e’’; yellow arrows). Altogether these results demonstrate that in contrast to its protein expression, *LoxL3* RNA is primarily expressed by bona fide MCT fibroblasts/FAPs some of which also express *MyoD1*.

## LPM-derived fibroblasts fuse to growing myofibers

The observation that *LoxL3* and *PDGFRα* expressing cells could also express myogenic markers (Fig. 2e), together with the results of the RNA velocity analysis (Fig. 1f), led us to test whether these cells are LPM-derived fibroblasts that have switched on a myogenic program. We took advantage of *Prx1^Cre^* ^20^, a LPM-specific Cre driver line previously demonstrated not to be expressed in myogenic cells^20–22^ and crossed it to the *Rosa26R^tdTomato^* reporter^23^ leading to tdTomato expressing LPM-derived cells. FISH analysis was carried out on cultured cells derived from limb muscles. We find that *MyoD1* RNA is expressed also in a subset of LPM-derived fibroblasts (Supp. Fig. 3a-a’’) and that tdTomato MHC expressing myotubes do form (Fig. 3a-a’’’).

**Figure 3.**
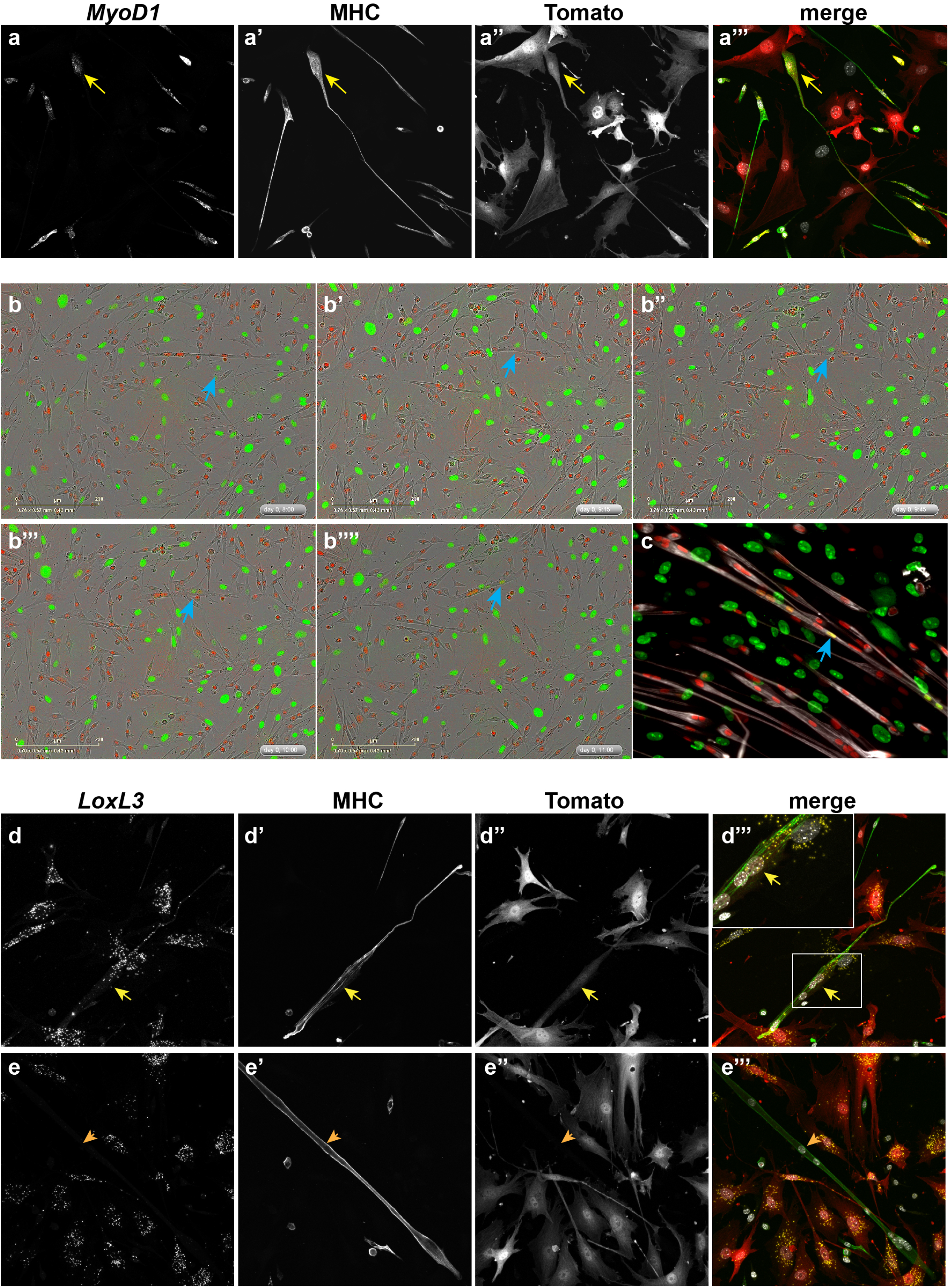
Cultured muscle interstitial cells fuse and contribute transcriptome to growing muscle fibers. Primary culture of myoblasts and interstitial cells derived from *Prx1^Cre^*; *Rosa^tdTomato^* following 24 hrs of differentiation demonstrates the presence of tdTomato expressing *MyoD1* (yellow) and MHC (green) expressing myofibers (a). Snapshots of live-cell imaging of *Prx1^Cre^*; *Rosa^nt-ng^*-derived primary culture after 48 hrs in differentiation media revealing fusion of fibroblast (blue arrow) into growing myofiber (b-b’’’’). Following 72 hrs in differentiation media, fibers marked by MHC (grey) with multiple fusion events are observed (c; blue arrows). FISH for *LoxL3* on primary culture of myoblasts and interstitial cells derived from *Prx1^Cre^*; *Rosa^tdTomato^* following 48 hrs of differentiation demonstrates that only in tdTomato-positive myofiber (d-d’’’) but not in tdTomato-negative (e-e’’’) fibers *LoxL3* RNA is observed.

The observation of tdTomato-expressing myotubes could be a consequence of several mechanisms that facilitate the introduction or transfer of the fibroblast-derived tdTomato RNA into the myofibers. Such mechanisms could include fusion of fibroblasts into the myofibers, extracellular vesicles released to the media that are then taken up by the myofibers or cellular extensions such as cytonemes between cell types^24–26^. The observation of adjacent *MyoD1* expressing cells, where one is tdTomato-positive and the other negative (Supp, Fig. 3a; yellow arrow) suggested this is not a random event and may result from a regulated process such as cell fusion, an event that is essential for myofiber growth. To directly test whether fibroblasts fuse into growing fibers we took advantage of the *Rosa^nt-ng^* reporter line in which all nuclei express tdTomato but upon Cre activity nuclear EGFP is switched on instead ^27^, thus enabling tracking of the distinct nuclei even in syncytia such as myofibers. Single cells were isolated from limb muscles of *Prx1^Cre^*; *Rosa^nt-ng^* P0 neonates and cultured in low serum to facilitate myogenic differentiation. In these cultures, all myogenic cells’ nuclei are tdTomato-expressing while all LPM-derived fibroblasts/FAPs’ nuclei express EGFP. Live cell imaging confirms EGFP expressing fibroblasts/FAPs fuse into growing myofibers (Fig. 3b-b’’’’; blue arrow and Supp. Movie1). Following 72 hours of in vitro differentiation ~7% of MHC expressing myotubes (n=276) harbor at least one EGFP expressing nucleus derived from such a fusion event (Fig. 3c).

That LPM-derived interstitial cells can fuse into growing myotubes raised the possibility that fibroblast-specific RNAs would be transferred along thus facilitating expression of fibroblast/FAP-associated genes such as *LoxL3* within the myotube. Accordingly, we find that only in tdTomato-positive fibers (i.e. have a contribution of at least one fibroblast) the fibroblast/FAP-specific RNAs of *LoxL3* and *PDGFRα* are found and actively transcribed as monitored by two fluorescent foci inside the nucleus (Fig. 3d-e’’’ and Supp. Fig. 3b-c’’’). Overall, these results reveal that not only do LPM-derived interstitial cells fuse into growing fibers but that they also maintain, at least to some extent, their initial identities’ transcriptional program.

A series of experiments marking proliferating cells, have demonstrated progenitors fuse to elongating myofibers primarily along the tips adjacent to the MTJs^5–7^. Although the previous experiments did not monitor the identity of these progenitors, *Prx1^Cre^; Rosa^nt-ng^* enable dissection of whether some of the fused cells are LPM-derived fibroblasts. *Prx1^Cre^; Rosa^nt-ng^* mice demonstrate that, as expected, the myofibers’ nuclei are primarily tdTomato-positive while EGFP expressing nuclei are found along the MTJs and in between myofibers (Fig. 4a). Should fibroblasts fuse into the growing myofibers also in vivo, we would expect that in proximity to the myofiber tips, adjacent to the MTJs, EGFP-expressing nuclei will be found. IMARIS 3D analysis of confocal images, demonstrates this is the case. Notably, these EGFP nuclei are primarily located close to the MTJs (Fig. 4a-b; Supp. Fig. 4a-a’; Supp. Movie2). Altogether these results demonstrate that fibroblast fusion into myofibers occurs as part of the normal developmental program and confirm the in vitro observations of the dual origin of cells that make up the myofiber.

**Figure 4.**
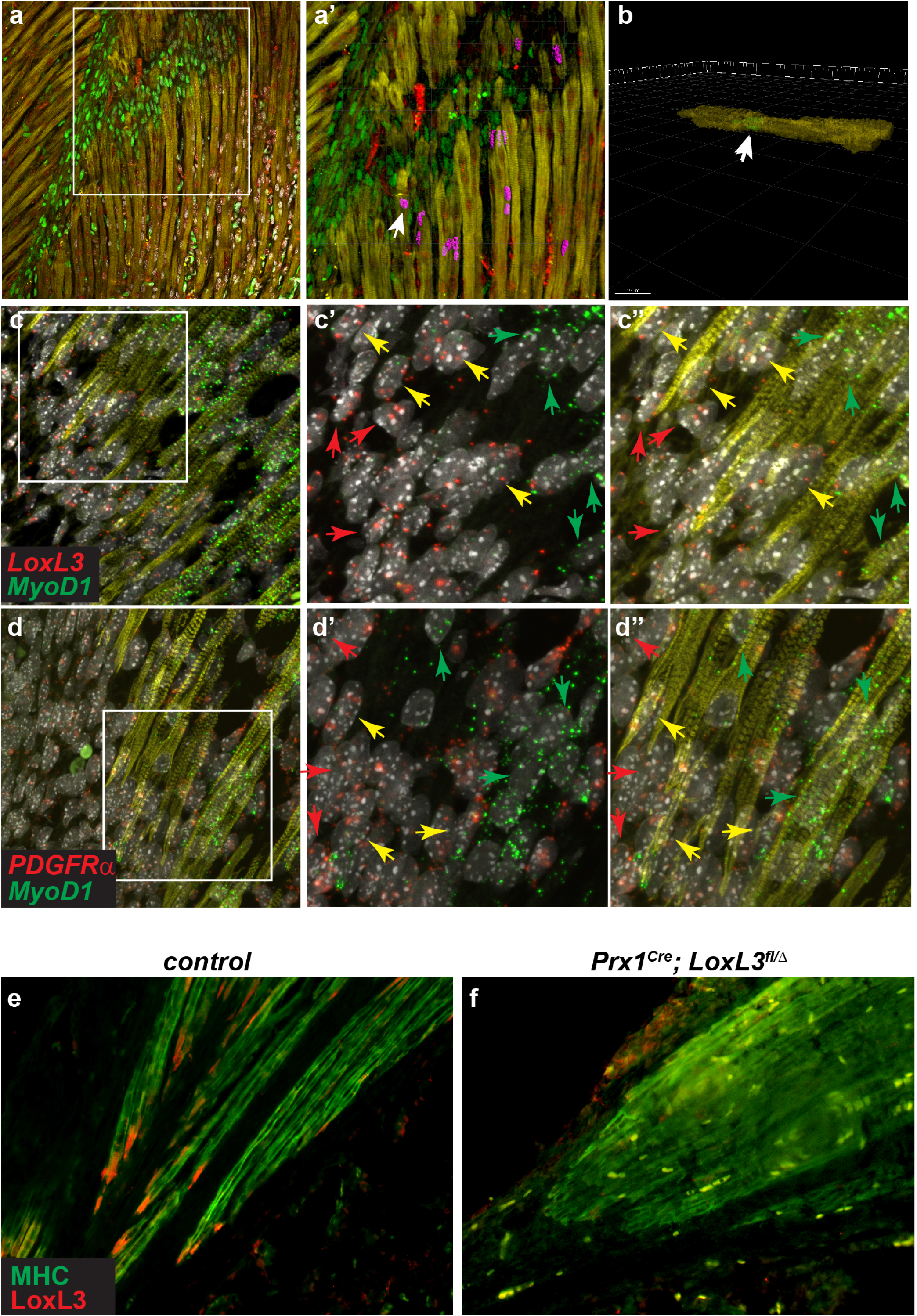
LPM-derived cells fuse into myofibers along the MTJ. P0 *Prx1^Cre^*; *Rosa^nt-ng^* limbs were subjected to immunostaining for MHC (yellow), EGFP (green) and tdTomato (red) (a) followed by IMARIS analysis demonstrates LPM-derived cells fuse into fibers at the proximity of the MTJ (a’, purple; b-lateral view of fiber section marked by white arrow in a’) (a,b). FISH for *LoxL3* and *MyoD1* (c-c’’) or *PDGFRα* and *MyoD1* (d-d’’) on E15.5 limb sections. Tagged images show magnification of boxed areas in c and d. Staining demonstrates spatial localization of the distinct nuclei at the fiber tips those located towards the center of the muscle express only *MyoD1* (green arrows in c’, d’), those closer to the tips (but also outside) express *MyoD1* and fibroblastic markers (yellow arrows in c’, d’) and those outside of the fiber towards the tendon express only fibroblastic markers (red arrows in c’, d’). Immunostaining for MHC (green) and LoxL3 (red) on E17.5 *control* (e) and *Prx1^Cre^LoxL3^fl/∆^*(f) demonstrates LoxL3 is missing at myofiber tips following its deletion in fibroblasts.

Fusion between myoblasts has been shown to be dependent on the transmembrane fusogene *myomaker* (*Mymk*)^28^. To test whether the LPM-derived cells fuse into the myofiber in a *Mymk*-dependent manner we deleted *Mymk* in the LPM lineage using the *Prx1^Cre^*, however no MTJ abnormalities were observed (Supp. Fig. 4b,c). Overall, these observations suggest that the fusion of LPM-derived cells into the myofiber is dependent on a mechanism not involving *Mymk* in fibroblasts and potentially consistent with the lack of need for *Mymk* in myofibers for fusion ^29,30^.

The finding that LPM-derived fibroblasts fuse into the myofiber in proximity to the MTJs led us to test whether *LoxL3*- and *PDGFRα*-expressing nuclei can be found within myofibers adjacent to the tips. FISH data for these fibroblast markers and for *MyoD1*, establishes that adjacent to the myofiber tips, both within them but also outside, nuclei actively transcribing fibroblastic markers and *MyoD1* are found [Fig. 4c-d (yellow arrows in c’ and d’); Supp. Fig. 4d-f’’’]. Interestingly, the nuclei located towards the center of the fiber express only *MyoD1* (green arrow in Fig. 4c’ and d’) whereas those located farther away and outside of the fiber express only the fibroblastic marker (red arrow in Fig. 4c’ and d’). Since some of the nuclei expressing both myogenic and fibrogenic markers are located at the tip of the fiber where MHC expression is sometimes less prominent, we were not able to conclusively quantify the number of these nuclei that are fully contained within the fiber (Supp. Fig. 4g).

In accordance with the above results, *LoxL3* deletion in the LPM-derived cells using the *Prx1^Cre^* deleter line but not in the myogenic progenitors using *Pax3^Cre^* line, results in MTJ defects (Supp. Fig. 4h,i) and in loss of LoxL3 expression from the myofiber tips (Fig. 4e,f). Overall, these results demonstrate that the myofiber tips at the MTJ region encompass unique nuclei that are of LPM-origin that also express fibroblastic markers and that this hybrid nature of the muscle fiber is critical for formation and maintenance of the MTJs.

## Discussion

Here we explored the mechanisms underlying the building of the musculoskeletal system focusing on the MTJs. Single cell resolution techniques allowed us to revisit the paradigms depicting muscle development. We identified, for the first time, that LPM-derived fibroblasts transdifferentiate by switching on the myogenic program and fuse into the myofiber. We demonstrate this fusion is part of the normal developmental program generating syncytial myofibers with hybrid developmental origins. Our results suggest this process is essential for MTJ formation. Reinforcing our observations, a unique group of nuclei expressing myogenic and fibrogenic genes (MTJ-B), the majority of which are shared by the dual identity cells, located at the MTJs, was recently identified in single nuclei sequencing of adult and regenerating myofibers ^31^ further suggesting the hybrid nature of muscle fibers.

Recent work has demonstrated the tendon-bone junctions is also composed of two cell types – tendon fibroblasts and chondrocytes^32,33^. Interestingly, this junction is dependent on TGFβ and BMP4 signaling^17,32,33^. Notably, tendon development is highly dependent on TGFβ signaling^34,35^ and BMP4 activity is prominent in muscle fiber tips^11^. Our results also point to an involvement of these two pathways in the dual identity cells that contribute to the MTJs putting forward the possibility that cells with dual programs are required for generating junctions between tissues.

## Acknowledgements

We are grateful to Dr. E. Suss-Toby and M. Holdengreber from the BCF Bioimaging Center (Faculty of Medicine, Technion); to L. Linde and R. Modai-Hod from the BCF Genomics Center (Faculty of Medicine, Technion) for bioinformatic analyses to members of the animal facility for excellent technical assistance. We further thank T. Schultheiss for critical reading of the manuscript and for fruitful discussions. P.H. was supported by grants from the Israeli Science Foundation (1072/13) and (1111/18), by A.D.I. (Association Duchenne Israel) and by the Rappaport Family Institute. The work in the laboratory of D.P.M. is supported by grants from the National Institutes of Health (R01AR068286, R01AG059605) and Pew Charitable Trusts.

## Author contributions

W.Y.B. conducted the majority of the experiments. O.K.S participated in analyzing the *LoxL3* MTJ phenotypes. S.Z.E. and S.M. assisted in cell culture assays. C.S and D.P.M. participated in the *myomaker* experiments. W.Y.B. and P.H planned the study, supervised and analyzed the data. D.P.M. and P.H. wrote the manuscript.

## METHODS

### Mice

All experiments involving mice conform to the relevant regulatory standards (institutional and national animal welfare laws, guidelines and policies). Embryonic day (E) was staged according to Kaufmann (^36^); noon of the day a vaginal plug was observed was marked as E0.5.

The *LoxL3* allele and the *Prx1^Cre^* were previously described^12,37^, respectively. *Rosa26R^tdTomato^* (also known as Ai9) ^23^ and *Rosa26^nTnG^* were purchased from JAX mice.

### Mouse Genotyping

Mice were PCR genotyped using the following primers:

*LoxL3*: LoxL3-5armWTF: GCCAGGGTGAAGTGAAAGAC; LoxL3-CritWTR: GATCTGGGATGCTGAAGACC; Tm1a-5mut-R1: GAACTTCGGAATAGGAACTTCG. 300 and 100 bp represent wild-type and mutant PCR products, respectively.

Cre: CreORF REV: ATCCAGGTTACGGATATAGT; CreORF FWD: ATCCGAAAAGAAAACGTTGA. 500 bp represent a cre positive PCR product.

nTnG: nT-nG_1: CCAGGCGGGCCATTTACCGTAAG; nT-nG_2: GGAGCGGGAGAAATGGATATG; nTnG 3: AAAGTCGCTCTGAGTTGTTAT. 603 and 320 bp represent wild-type and homozygous presence of nTnG PCR products, respectively.

tdTomato: RosaTomato WT_FWD: AAGGGAGCTGCAGTGGAGTA; RosaTomato WT_REV: CCGAAAATCTGTGGGAAGTC. Fragment size of 196 bp represent the tomato transgene, while a fragment size of 297 bp represent the wild-type.

### Fluorescent in situ hybridization (FISH)

FISH was carried out using the RNAScope system according to the manufacturers’ protocol^38^, on FFPE forelimb and hindlimb sections and also in primary cell cultures. Briefly, the sections were baked in a dry oven for 1 hr at 60°C, deparaffinized using xylene and ethanol, air dried and incubated in hydrogen peroxide for 10 min. For the antigen retrieval step, slides were boiled in an anti-retrieval solution for 2 min, transferred to ethanol for 3 min and air dried. Slides were then incubated with an RNAScope special protease for 30 min. Designed target probes of *LoxL3* (431359; RNAScope), *MyoD1* (316081-C2; RNAScope), or *PDGFRα* (480661-C3; RNAScope) were added to the slides and incubated for 2 hr at 40°C. A series of probe amplifiers were added afterwards. At the end of the protocol, after adding the fluorophores, immunofluorescent staining was carried out. For cultured adherent cells, a similar protocol was carried out with the exception that the hydrogen peroxide incubation commenced directly after fixation (4% PFA).

### Immunohistochemistry

Section and whole-mount immunohistochemistry were performed essentially as previously described^39^. The following antibodies were used: anti-Myosin (A4.1025, 1:300; DSHB); anti-GFP (A6455, 1:500; Invitrogen); anti-RFP (5F8, 1:500; Chromotek); anti-LOXL3^12^ (1:100).

### Generation of primary cultures

P0 embryo limb muscles were sliced into pieces, incubated with collagenase for 1 hour and then with trypsin for half an hour. Tissues were then centrifuged for 15 minutes at 1300 rpm, resuspended in DMEM (containing 1% L-glutamine and 1% penicillin and streptomycin (PS) + 10% FCS, filtered, re-centrifuged, re-suspended in BIO-AMF-2 (Biological Industries) and plated on gelatin coated plates. Differentiation was induced by incubating the cells in DMEM (containing 1% L-glutamine and 1% PS) + 4% horse serum for 3 days.

### 10x Genomics Single Cell General Cell Preparation

Single cell separation and library construction according to 10x protocol (Chromium Single Cell 3’ Library & Gel Bead Kit v2). Briefly, the MTJ region was dissected from P0 neonates and single cells were isolated (as described) and suspended with BIO-AMF-2 (Biological Industries). The cells were mixed thoroughly using a wide-bore pipette tip and counted. Cells were centrifuged at 1300 rpm for 5 min at RT, suspended with 0.04% BSA PBS, re-centrifuged and then re-suspended with the appropriate volume of 0.04% BSA PBS to achieve a target cell concentration in the range of 700-1200 cell/ul. Following cell isolation, library construction was immediately carried out. Cell Ranger software was used for data QC and extraction of transcripts’ counts from raw data.

### Bioinformatic analysis

Seurat R package was used for filtering, clustering and expression distribution of selected cluster-specific genes. SingleR R Package was used for unbiased cell type recognition of scRNA-seq. Cells with the following parameters were excluded: >8% mitochondrial UMI counts; less than 200 unique gene counts; over 4,000 unique gene counts. Overall, 10,456 cells entered the analysis and 9,238 cells were used for the bioinformatics analysis after filtration. In addition, genes detected in less than 3 cells were filtered out. WebGestalt (WEB-based Gene SeT AnaLysis Toolkit) a functional enrichment analysis web tool and Ingenuity Pathway Analysis (Qiagen) were used to identify specific pathways and processes within the data.

RNA velocity analysis was conducted as follows: A loom file from 10x output using velocyto.R with default parameters was used^19^. In summary estimating RNA Velocity using Seurat was created and imported to R and an RNA velocity analysis was conducted for the whole data. The cluster labels from the initial analysis were assigned to the velocity analysis in order to compare between the results. Cells that didn’t assign to any initial cluster were assigned as cluster “0”. RNA velocity was calculated on this data. Furthermore, cells that were labeled as fibroblasts, myoblasts, satellite cells, myocytes and dual identity were extracted and analyzed using RNA velocity without re-clustering the data.

## References

1 Kieny, M. & Chevallier, A. Autonomy of tendon development in the embryonic chick wing. J Embryol Exp Morphol 49, 153–165 (1979).

2 Schweitzer, R. et al. Analysis of the tendon cell fate using Scleraxis, a specific marker for tendons and ligaments. Development (Cambridge, England) 128, 3855–3866 (2001).

3 Tajbakhsh, S. & Buckingham, M. The birth of muscle progenitor cells in the mouse: spatiotemporal considerations. Curr Top Dev Biol 48, 225–268, doi:10.1016/s0070-2153(08)60758-9 (2000).

4 Wachtler, F., Christ, B. & Jacob, H. J. On the determination of mesodermal tissues in the avian embryonic wing bud. Anat Embryol (Berl) 161, 283–289, doi:10.1007/BF00301826 (1981).

5 Gu, J. M. et al. An NF-kappaB--EphrinA5-Dependent Communication between NG2(+) Interstitial Cells and Myoblasts Promotes Muscle Growth in Neonates. Developmental cell 36, 215–224, doi:10.1016/j.devcel.2015.12.018 (2016).

6 Kitiyakara, A. & Angevine, D. M. A Study of the Pattern of Postembryonic Growth of M. Gracilis in Mice. Developmental biology 8, 322–340, doi:10.1016/0012-1606(63)90033-2 (1963).

7 Williams, P. E. & Goldspink, G. Longitudinal growth of striated muscle fibres. J Cell Sci 9, 751–767 (1971).

8 Tsujimura, T., Kinoshita, M. & Abe, M. Response of rabbit skeletal muscle to tibial lengthening. Journal of orthopaedic science: official journal of the Japanese Orthopaedic Association 11, 185–190, doi:10.1007/s00776-005-0991-8 (2006).

9 Zhang, M. & McLennan, I. S. During secondary myotube formation, primary myotubes preferentially absorb new nuclei at their ends. Dev Dyn 204, 168–177, doi:10.1002/aja.1002040207 (1995).

10 Edom-Vovard, F., Bonnin, M. A. & Duprez, D. Misexpression of Fgf-4 in the chick limb inhibits myogenesis by down-regulating Frek expression. Developmental biology 233, 56–71, doi:10.1006/dbio.2001.0221 (2001).

11 Wang, H. et al. Bmp signaling at the tips of skeletal muscles regulates the number of fetal muscle progenitors and satellite cells during development. Developmental cell 18, 643–654, doi:10.1016/j.devcel.2010.02.008 (2010).

12 Kraft-Sheleg, O. et al. Localized LoxL3-Dependent Fibronectin Oxidation Regulates Myofiber Stretch and Integrin-Mediated Adhesion. Developmental cell 36, 550–561, doi:10.1016/j.devcel.2016.02.009 (2016).

13 Baumeister, A., Arber, S. & Caroni, P. Accumulation of muscle ankyrin repeat protein transcript reveals local activation of primary myotube endcompartments during muscle morphogenesis. The Journal of cell biology 139, 1231–1242, doi:10.1083/jcb.139.5.1231 (1997).

14 Esteves de Lima, J., Bonnin, M. A., Birchmeier, C. & Duprez, D. Muscle contraction is required to maintain the pool of muscle progenitors via YAP and NOTCH during fetal myogenesis. eLife 5, doi:10.7554/eLife.15593 (2016).

15 Jo, C. H., Lim, H. J. & Yoon, K. S. Characterization of Tendon-Specific Markers in Various Human Tissues, Tenocytes and Mesenchymal Stem Cells. Tissue engineering and regenerative medicine 16, 151–159, doi:10.1007/s13770-019-00182-2 (2019).

16 Subramanian, A. & Schilling, T. F. Thrombospondin-4 controls matrix assembly during development and repair of myotendinous junctions. eLife 3, doi:10.7554/eLife.02372 (2014).

17 Maeda, T. et al. Conversion of mechanical force into TGF-beta-mediated biochemical signals. Curr Biol 21, 933–941, doi:10.1016/j.cub.2011.04.0070960-9822(11)00423-4 [pii] (2011).

18 Wang, J., Duncan, D., Shi, Z. & Zhang, B. WEB-based GEne SeT AnaLysis Toolkit (WebGestalt): update 2013. Nucleic Acids Res 41, W77–83, doi:10.1093/nar/gkt439 (2013).

19 La Manno, G. et al. RNA velocity of single cells. Nature 560, 494–498, doi:10.1038/s41586-018-0414-6 (2018).

20 Logan, M. et al. Expression of Cre Recombinase in the developing mouse limb bud driven by a Prxl enhancer. Genesis 33, 77–80 (2002).

21 Colasanto, M. P. et al. Development of a subset of forelimb muscles and their attachment sites requires the ulnar-mammary syndrome gene Tbx3. Dis Model Mech 9, 1257–1269, doi:10.1242/dmm.025874 (2016).

22 Seo, H. S. & Serra, R. Deletion of Tgfbr2 in Prx1-cre expressing mesenchyme results in defects in development of the long bones and joints. Developmental biology 310, 304–316, doi:10.1016/j.ydbio.2007.07.040 (2007).

23 Madisen, L. et al. A robust and high-throughput Cre reporting and characterization system for the whole mouse brain. Nature neuroscience 13, 133–140, doi:10.1038/nn.2467 (2010).

24 Sampath, S. C., Sampath, S. C. & Millay, D. P. Myoblast fusion confusion: the resolution begins. Skelet Muscle 8, 3, doi:10.1186/s13395-017-0149-3 (2018).

25 van Niel, G., D’Angelo, G. & Raposo, G. Shedding light on the cell biology of extracellular vesicles. Nature reviews. Molecular cell biology 19, 213–228, doi:10.1038/nrm.2017.125 (2018).

26 Yamashita, Y. M., Inaba, M. & Buszczak, M. Specialized Intercellular Communications via Cytonemes and Nanotubes. Annual review of cell and developmental biology 34, 59–84, doi:10.1146/annurev-cellbio-100617-062932 (2018).

27 Prigge, J. R. et al. Nuclear double-fluorescent reporter for in vivo and ex vivo analyses of biological transitions in mouse nuclei. Mammalian genome: official journal of the International Mammalian Genome Society, doi:10.1007/s00335-013-9469-8 (2013).

28 Millay, D. P. et al. Myomaker is a membrane activator of myoblast fusion and muscle formation. Nature 499, 301–305, doi:10.1038/nature12343 (2013).

29 Goh, Q. & Millay, D. P. Requirement of myomaker-mediated stem cell fusion for skeletal muscle hypertrophy. eLife 6, doi:10.7554/eLife.20007 (2017).

30 Petrany, M. J., Song, T., Sadayappan, S. & Millay, D. P. Myocyte-derived Myomaker expression is required for regenerative fusion but exacerbates membrane instability in dystrophic myofibers. JCI insight 5, doi:10.1172/jci.insight.136095 (2020).

31 Kim, M. et al. Single-nucleus transcriptomics reveals functional compartmentalization in syncytial skeletal muscle cells. bioRxiv, 2020.2004.2014.041665, doi:10.1101/2020.04.14.041665 (2020).

32 Blitz, E., Sharir, A., Akiyama, H. & Zelzer, E. Tendon-bone attachment unit is formed modularly by a distinct pool of Scx- and Sox9-positive progenitors. Development (Cambridge, England) 140, 2680–2690, doi:10.1242/dev.093906 (2013).

33 Zelzer, E., Blitz, E., Killian, M. L. & Thomopoulos, S. Tendon-to-bone attachment: from development to maturity. Birth defects research. Part C, Embryo today: reviews 102, 101–112, doi:10.1002/bdrc.21056 (2014).

34 Havis, E. et al. Transcriptomic analysis of mouse limb tendon cells during development. Development (Cambridge, England) 141, 3683–3696, doi:10.1242/dev.108654 (2014).

35 Pryce, B. A. et al. Recruitment and maintenance of tendon progenitors by TGFbeta signaling are essential for tendon formation. Development (Cambridge, England) 136, 1351–1361, doi:10.1242/dev.027342 (2009).

36 Kaufmann, M. H. The Atlas of Mouse Development. (1992).

37 Logan, M. & Tabin, C. Targeted gene misexpression in chick limb buds using avian replication-competent retroviruses. Methods (San Diego, Calif 14, 407–420 (1998).

38 Wang, F. et al. RNAscope: a novel in situ RNA analysis platform for formalin-fixed, paraffin-embedded tissues. The Journal of molecular diagnostics: JMD 14, 22–29, doi:10.1016/j.jmoldx.2011.08.002 (2012).

39 Hasson, P. et al. Tbx4 and Tbx5 acting in connective tissue are required for limb muscle and tendon patterning. Developmental cell 18, 148–156 (2010).

