## Supplemental figures for "Fibroblast fusion to the muscle fiber regulates myotendinous junction formation"

**Supp. Fig. 1. scRNA seq highlights dual characteristics of cluster 19 cells.** Violin plots depicting cell markers of myocytes (a), tenocytes (b), endothelial cells (c), smooth muscle cells (d) myogenic cells (e) and fibroblasts (f). Venn diagram depicting differentially expressed gene comparisons between myogenic and fibroblastic cell clusters and with dual identity cluster (g). UMAP plot derived from RNA velocity analysis of all clusters (h). Violin plot of *LoxL3* expression in distinct clusters (i).

**Supp. Fig. 2. LoxL3 expressed in myofiber tips is required for MTJs.** Wholemount MHC immunostaining on E18.5 forelimbs of LoxL3 mutant (a, a'), control (b) and LoxL3 mutant (c) embryos. Immunostaining for LoxL3 (green) and MHC (red) on E12.5 forelimb (d, d'), E13.5 rib (e) and E14.5 (f), E16.5 (g-i) forelimbs demonstrates LoxL3 marks muscle tips at multiple stages and in distinct regions. Immunostaining for LoxL3 (green) and Laminin (red) on E16.5 sections demonstrates LoxL3 is expressed inside the myofibers (j, arrows). In *LoxL3* mutant embryos, no LoxL3 staining is observed demonstrating antibody specificity (k-k''). FISH staining for *PDGFRα* (l, l'' green) and *MyoD1* (l', l'' red) along with immunostaining for MHC (l'' grey) on primary cultured cells following 48 hrs of differentiation demonstrates the two markers highlight distinct cells.

**Supp. Fig. 3. LPM-derived cells contribute to myofibers.** FISH for *MyoD1* (a, a'' green) on primary cells derived from *Prx1$^{Cre}$; Rosa26$^{tdTomato}$* following 48hrs of differentiation. Yellow arrow marks two adjacent *MyoD1* expressing cells, one tdTomato positive and the other tdTomato negative. Orange arrowhead marks *MyoD1* expressing cell that is tdTomato negative. FISH for *PDGFRα* on primary culture of myoblasts and interstitial cells derived from *Prx1$^{Cre}$*; *Rosa$^{tdTomato}$* following 48 hours of differentiation demonstrates that only in tdTomato-positive myofiber (b-b''') but not in tdTomato-negative (c-c''') fibers *PDGFRα* RNA is observed.

**Supp. Fig. 4. Nuclei at myofiber tips express interstitial markers.** Immunostaining for MHC (yellow), GFP and Tomato on a P0 *Prx1$^{Cre}$; Rosa26$^{nt\text{-}ng}$* limb (a) demonstrating process of IMARIS analysis to identify LPM-derived GFP positive cells inside myofibers (a' purple). MHC wholemount immunostaining of E16.5 *myomaker$^{fl/+}$* (b) or *Prx1$^{Cre}$; myomaker$^{fl/fl}$* (c) limbs. MHC immunostaining (yellow) and FISH for *LoxL3* (red) and *MyoD1* (green) (d), *LoxL3* (red) and *PDGFRα* (blue) (e), *MyoD1* (green) and *PDGFRα* (blue) (f). Tagged images show higher magnification of boxed regions in non-tagged ones (d-f'''). High magnification of myofibers tips (MHC, green) along with FISH for *LoxL3* (red) and *MyoD1* (yellow) (g). White arrows denote examples of *LoxL3* expressing nuclei that we are not able to determine if located in or out of myofiber. MHC immunostaining on (E16.5) control (h) and *Prx1*$^{Cre}$*LoxL3*$^{fl/\Delta}$ (i) demonstrates MTJ defects are observed following *LoxL3* deletion in LPM (black arrow).

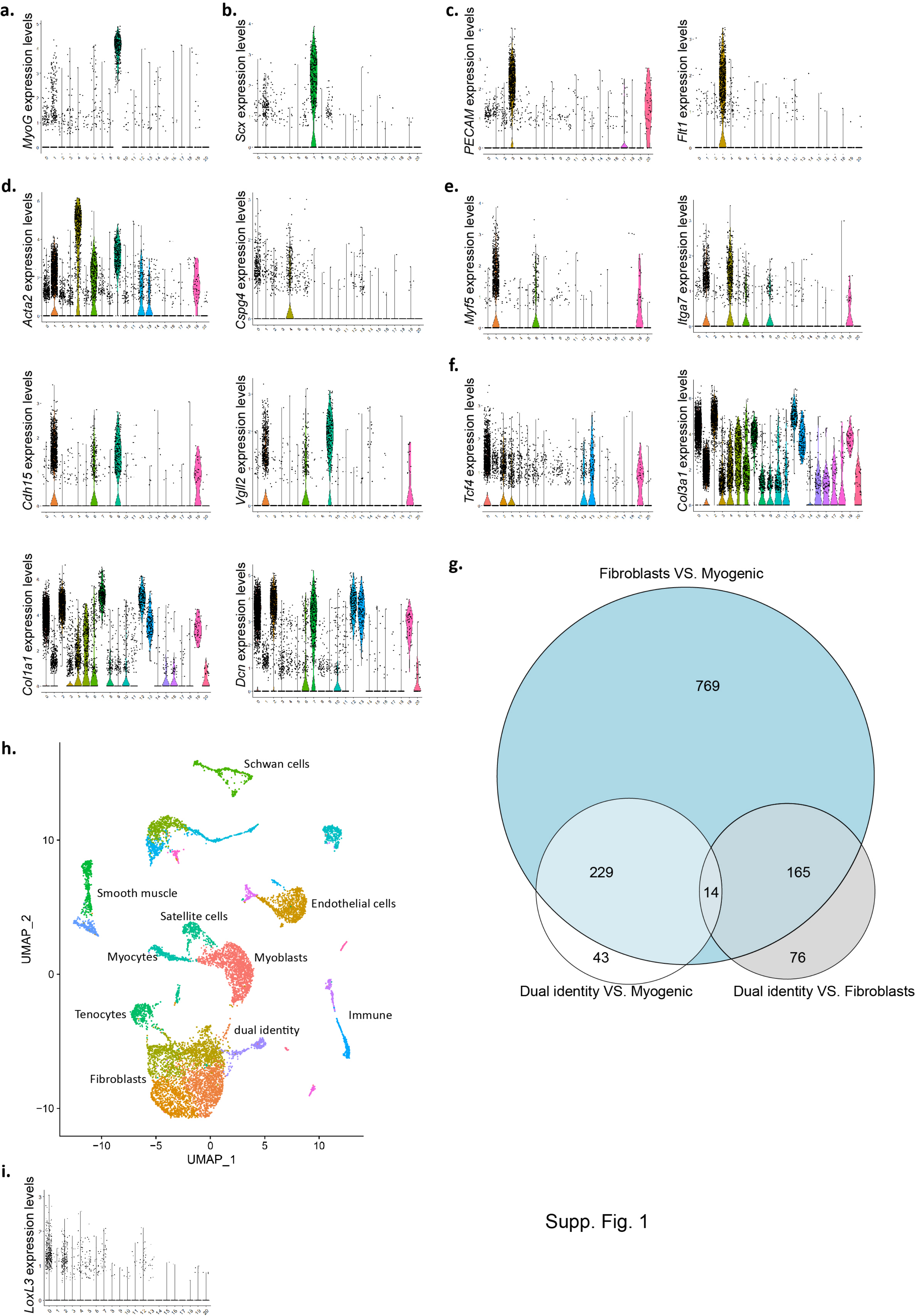
a.
MyoG expression levels
b.
Scx expression levels
c.
PECAM expression levels
Flt1 expression levels
d.
Acta2 expression levels
Cspg4 expression levels
e.
Myf5 expression levels
Itga7 expression levels
Cdh15 expression levels
Vgll2 expression levels
f.
Tcf4 expression levels
Col3a1 expression levels
Col1a1 expression levels
Dcn expression levels
g.
Fibroblasts VS. Myogenic
769
229
14
165
43
76
Dual identity VS. Myogenic
Dual identity VS. Fibroblasts
h.
Schwan cells
Smooth muscle
Endothelial cells
Satellite cells
Myocytes
Myoblasts
Tenocytes
dual identity
Immune
Fibroblasts
UMAP_2
UMAP_1
i.
LoxL3 expression levels

Supp. Fig. 1

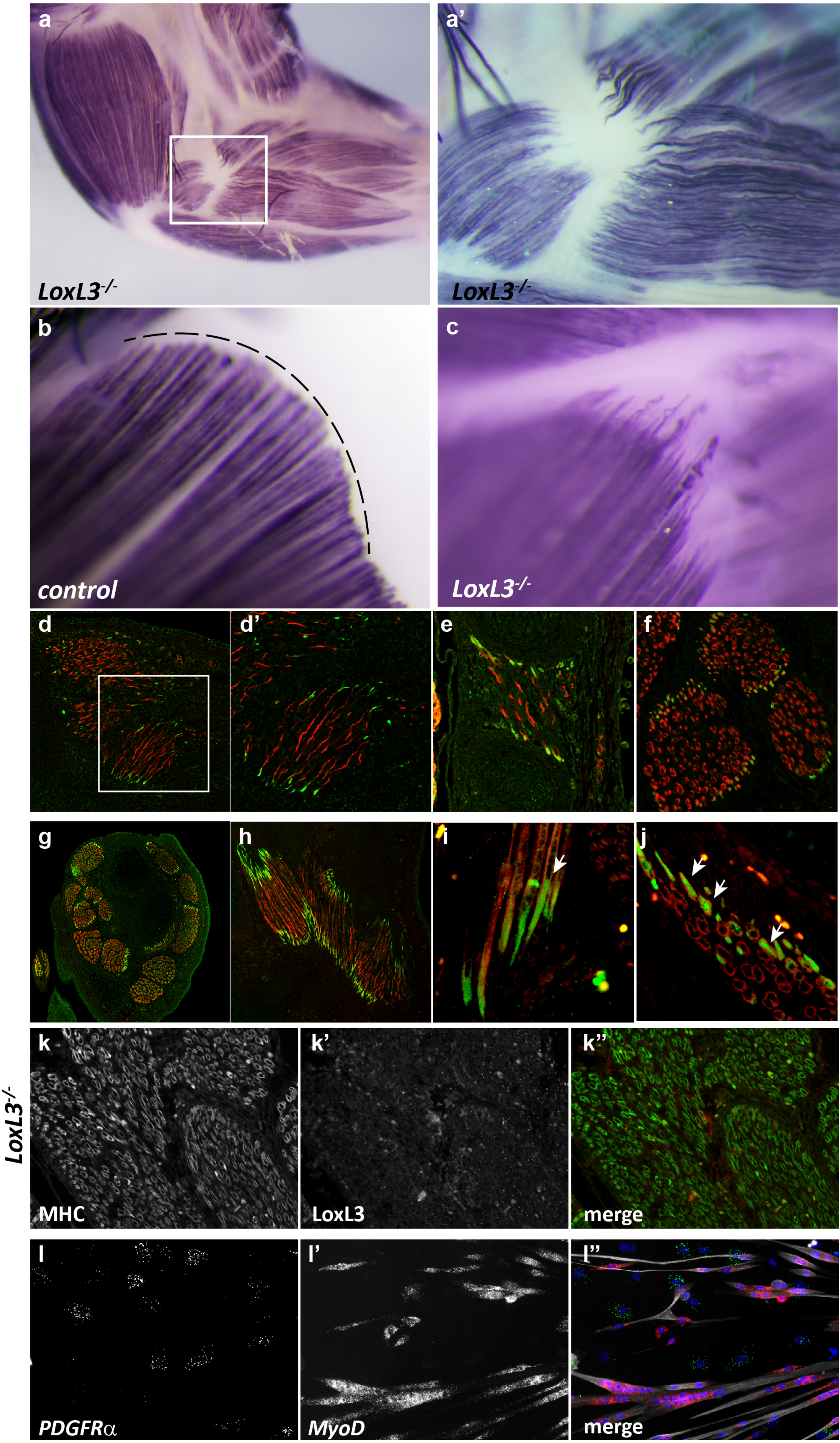
a
LoxL3-/-
a'
LoxL3-/-
b
control
c
LoxL3-/-
d
d'
e
f
g
h
i
j
k
k'
k''
LoxL3-/-
MHC
LoxL3
merge
l
l'
l''
PDGFRα
MyoD
merge

Supp. Fig. 2

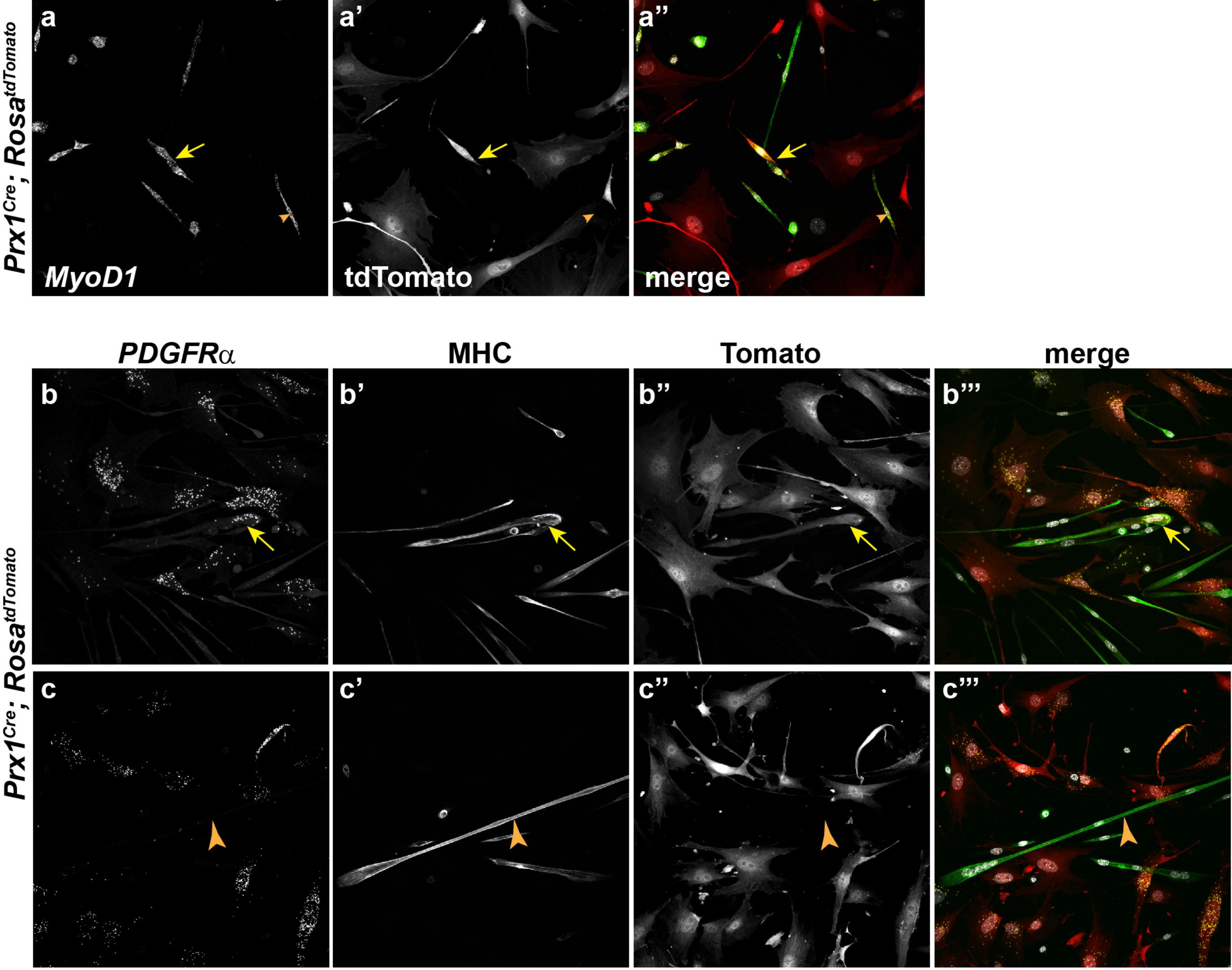


**Supp. Figure 3**

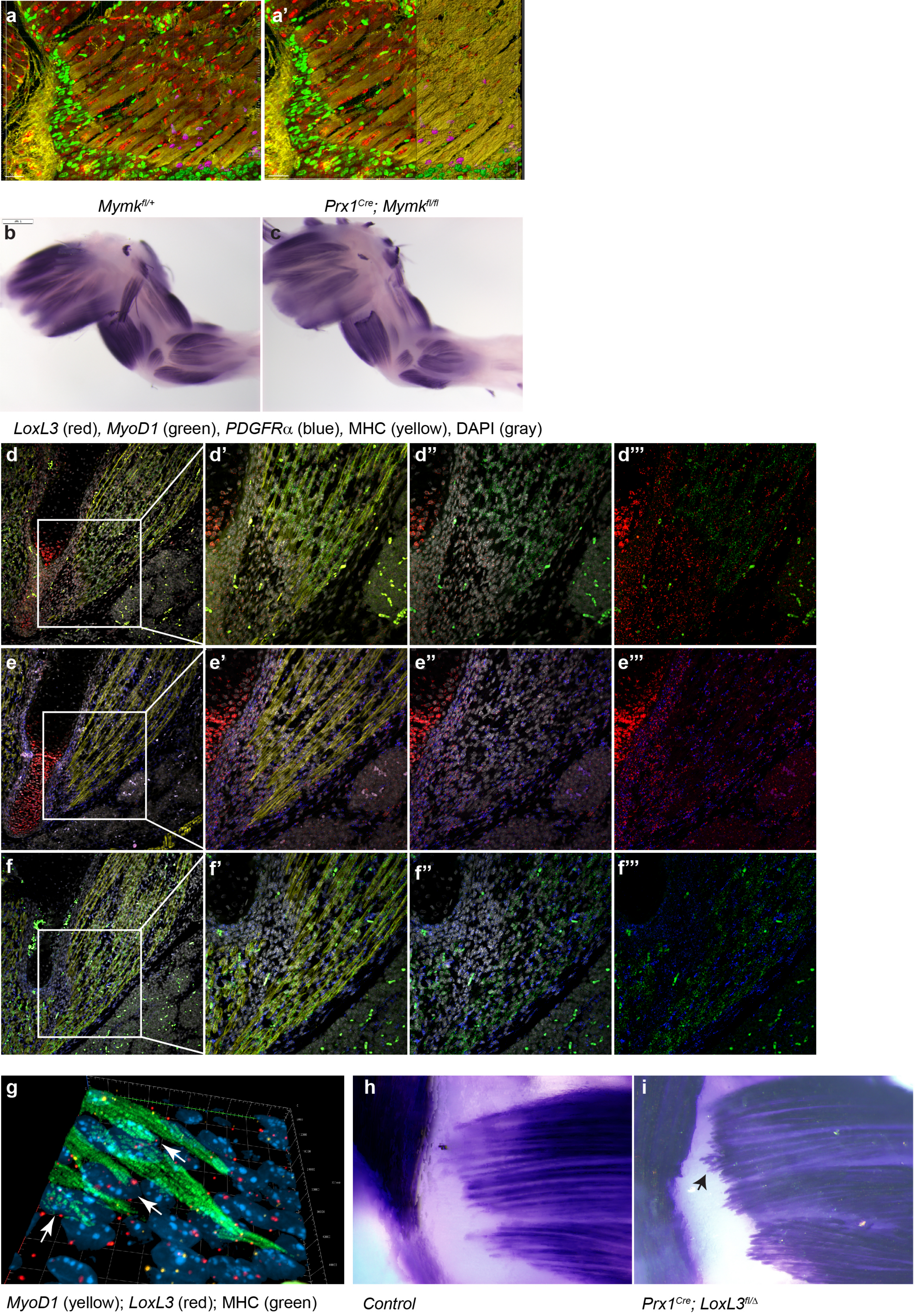


**Supp. Figure 4**
